# ATAC-seq signal processing and recurrent neural networks can identify RNA polymerase activity

**DOI:** 10.1101/531517

**Authors:** Ignacio J. Tripodi, Murad Chowdhury, Robin Dowell

## Abstract

Nascent transcription assays are the current gold standard for identifying regions of active transcription, including markers for functional transcription factor (TF) binding. Here we present a signal processing-based model to determine regions of active transcription genome-wide using the simpler assay for transposase-accessible chromatin, followed by high-throughput sequencing (ATAC-seq). The focus of this study is twofold: First, we perform a frequency space analysis of the “signal” generated from ATAC-seq experiments’ short reads, at a single-nucleotide resolution, using a discrete wavelet transform. Second, we explore different uses of neural networks to combine this signal with its underlying genome sequence in order to classify ATAC-seq peaks on the presence or absence of bidirectional transcription. We analyze the performance of different data encoding schemes and machine learning architectures, and show how a hybrid signal/sequence representation classified using recurrent neural networks (RNNs) yields the best performance across different cell types.

## Introduction

The study of transcription factor (TF) activity is key to understanding transcription regulatory networks. Capturing functional binding is essential to understand mechanistic aspects of how a biological perturbation results in changes in transcription activity. While binding detection assays such as ChIP-seq report on associations between a TF and DNA, they are not necessarily informative on the TF’s functional activity of altering RNA polymerase behavior. Nascent transcription assays are more informative in this regard. The current state-of-the-art marker for functional TF binding to a regulatory region is co-occurrence with proximal transcription (Azofeifa *et al.,* 2018). Typically this transcription is bidirectional and roughly originates at the TF binding motif (Melgar *et al.,* 2011; Azofeifa *et al.,* 2018). It is thought that TFs recognize their motif and, when functional, recruit RNA polymerase which transcribes the immediately flanking regions. At enhancers, this gives rise to short unstable transcripts referred to as (eRNA). Thus, the TF is not only bound but also altering transcription.

Nascent transcription experiments are, however, quite laborious, expensive, and require a large number of cells. In contrast, the assay for transposase-accessible chromatin, followed by high-throughput sequencing (ATAC-seq) has rapidly gained popularity since its inception, due to its ease of execution, small cell count requirements, and short time expenditure. However, ATAC-seq measures chromatin accessibility, not RNA polymerase activity. Most sites of bidirectional transcription co-occur with open chromatin regions detectable by ATAC-seq (Fig. 1). Yet only a small fraction of open chromatin regions harbor RNA polymerase activity. Reasoning that the presence of RNA polymerase may itself alter chromatin state in some subtle fashion, we wondered whether ATAC-seq could be utilized to discriminate peaks that overlap RNA polymerase activity from other open chromatin regions unrelated to active transcription.

**Figure 1.**
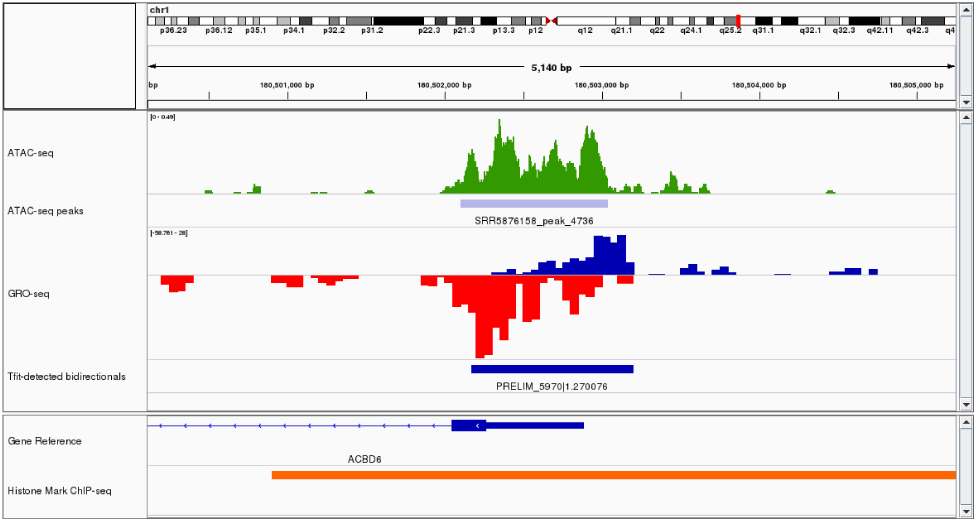
Integrative Genomics Viewer (IGV) snapshot depicting ATAC-seq signal (in green) (Kelso et al., 2017) and the associated peak call (light blue box) over a nascent transcription (Allen et al., 2014) (blue: positive strand transcription; red: negative strand transcription) bidirectional call (dark blue box) in cell type and condition matched data (chr1:180500000-180505000 in unperturbed HCT116). This region happens to correspond to an annotated gene promoter (ACBD6) and has an associated called H3K4me1 histone mark (orange box, ENCFF246BAX dataset).

Machine learning presents itself as a natural tool to classify data derived from genomics assays, particularly ATAC-seq. It has been utilized in a wide range of applications, from classifying types of chronic lymphocytic leukemia cells (Rendeiro *et al.,* 2016), to TF motif discovery (Setty and Leslie, 2015), or discriminating among brain cell types (Fullard *et al.,* 2018). Moreover, a recent study (Thibodeau *et al.,* 2018) attempted to identify gene enhancer regions using ATAC-seq peaks.

We approached the ATAC-seq peak classification problem as a signal processing task, taking the linearized chromosomal representation of mapped reads denoting OCRs as signal defining a peak. To this end we employed wavelet analysis, as it has been applied to different areas of genomics before, from predicting the position and shape of nucleosomes (Nguyen *et al.,* 2014) from histone marks, to detecting regions of ChIP-seq enrichment in less than favorable signal-to-noise ratio conditions (Mitra and Song, 2012), or extracting signatures from Hi-C correlation matrices (Chen *et al.,* 2016). Wavelet transforms extract the main frequency components from a signal, at a varying resolution for each component. They are ideal for discontinuous (choppy) signals, as is the case for ATAC-seq peaks, and for data compression applications. For a good introduction to wavelet analysis and common mother wavelet functions, see Graps et al (Graps, 1995).

We also explored different sequence representation schemes, reasoning that the positional sequence composition bias observed at enhancers and promoters (Azofeifa *et al.,* 2018) might improve our classifier. Convolutional neural networks apply small kernels in a sliding window across input sequences and have been shown to learn meaningful motif “features” from sequence data (Min *et al.,* 2017). In fact, deep learning sequence based techniques have been applied to a variety of tasks including prediction of enhancers (Min *et al.,* 2017), accessibility of DNA sequences (Kelley *et al.,* 2016), the effect of non-coding variants (Zhou and Troyanskaya, 2015), sequence specificity of binding proteins (Alipanahi *et al.,* 2015), and RNA protein coding potential (Hill *et al.,* 2018). Often in these models, nucleotides are represented using one-hot encoded vectors. Recurrent neural networks have also been shown to be able to learn from sequence data, owing to their ability to learn long term relationships and patterns. While it is possible to learn using the aforementioned one-hot encoding of input sequences, Hill et al (Hill *et al.,* 2018) found that use of an embedding input layer mapping individual nucleotides to a learned dense vector representation can lead to increased prediction performance. Furthermore, they found that certain nucleotides such as C and U, or C and G had lower cosine distances in the embedding vector space than between other pairs of nucleotides, showing that the learned embeddings held some biologically relevant meaning.

Ultimately, we combined these two sources of information into a signal/sequence hybrid, using recurrent neural networks to classify peaks. We rationalized this combination of underlying sequence with normalized read counts as a hybrid representation that “weights” each nucleotide by its context relevance (cell type, conditions, etc) from experimental data. Finally, while the focus of this study was on detecting signatures of RNA polymerase activity in ATAC-seq peaks, we also expanded the analysis to detecting histone marks and a combination of classification labels from both histone marks and nascent transcription.

## Methods

### Datasets

We utilized a collection of samples from different human cell lines and sources. ATAC-seq samples were selected from the public repositories based on the availability of both histone modification ChIP-seq data (specifically H3K4me1, H3K4me2, H3K4me3, H3K27ac, and H3K9ac) and nascent transcription data. In all cases, we matched cell type and sample conditions (untreated, or “DMSO”). Under this criteria, we obtained samples from myeloid B-cells (GM12878 (Schep *et al.,* 2015)), human embryonic stem cells (H1), colon carcinoma (HCT116 (Kelso *et al.,* 2017)) and leukemia lymphoblasts (K562 (Fuglerud *et al.,* 2017)). The samples retrieved from the Gene Expression Omnibus (GEO (Barrett *et al.,* 2013)) and ENCODE (The ENCODE Project Consortium, 2012) are listed on Supplemental Table 1 for ATAC-seq, Supplemental Table 2 for GRO-seq/PRO-seq, and Supplemental Table 3 for histone mark ChIP-seq.

The ATAC-seq samples were processed using the Nextflow-based (Di Tommaso *et al.,* 2017) ATACFlow (Tripodi *et al.,* 2018a) pipeline with default settings. The histone mark ChIP-seq peaks were downloaded from ENCODE (The ENCODE Project Consortium, 2012) using the combined replicates for each case. The nascent transcription datasets were processed using the Nextflow-based Nascent-Flow (Tripodi and Gruca, 2018) pipeline with default settings. Both Tfit (Azofeifa *et al.,* 2018) and FStitch (Azofeifa *et al.,* 2014) were then used to detect the bidirectional transcription regions, known to be regions of RNA polymerase loading and initiation. All samples and sequences were analyzed with respect to the GrCH38 human reference genome.

### Feature Engineering

We could separate the types of features derived from ATAC-seq peaks into three major categories: peak attributes, peak sequence, and peak signal. Normalized ATAC-seq peaks correspond to OCRs and are summarized into discrete peak signal features. To this end we utilized an evaluation window of 1kbp centered at each peak, resulting in 1000 features for each OCR. The signal features were thus derived from 1000 real values, representing the normalized number of reads in the evaluation window at single-nucleotide resolution.

### Signal Processing Encoding

We adopted a signal processing approach, speculating that there may be characteristic signatures in the peaks containing RNA polymerase (e.g. transcribed). The ATAC-seq peak “signal” itself (represented by the normalized read count on each nucleotide within a 1kbp window surrounding the peak’s midpoint) was examined in frequency space, using PyWavelets (Lee *et al.,* 2019). We performed a discrete wavelet transform (WL) on each ATAC-seq peak (Fig. 2), using the resulting frequency components (approximation coefficients) as features for a machine learning classifier. After a grid search for the optimal configuration, the “db38” wavelet function proved to be the most effective to classify ATAC-seq peaks, at one level of decomposition.

**Figure 2.**
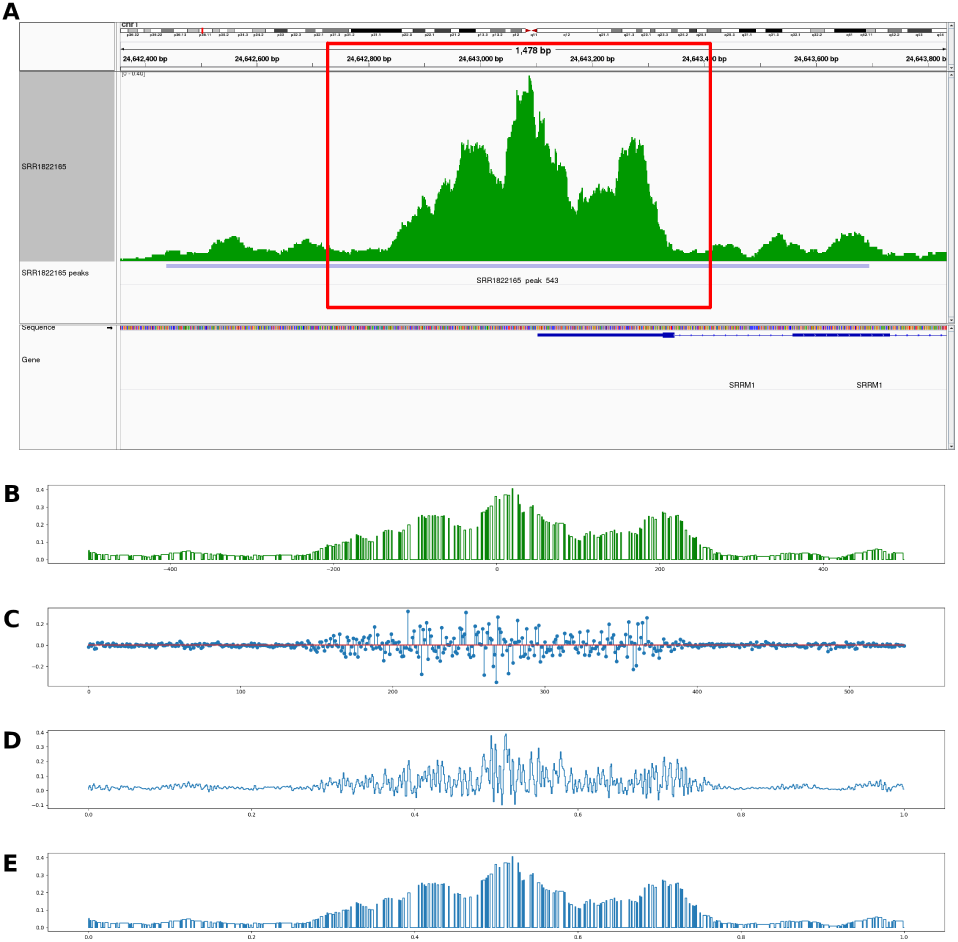
Example of wavelet analysis of an ATAC-seq peak. A A 1kb window (red box) around each ATAC-seq peak, as show in in the Integrative Genome Viewer (IGV) are analyzed (B). The level 1 approximation coefficients are shown in C, which are the features we use to train our machine learning model. For illustrative purposes, we show how the original signal (B) can be coarsely reconstructed by using only the level 1 coefficients (D), and a better reconstruction (E) is obtained using the detail coefficients.

As an alternative representation, we fed the 1000 features to an autoencoder (AE) layer consisting of a feed-forward artificial neural network with a single hidden layer. In this manner we reduce the number of dimensions while preserving an encoding of most of the important signal features. The encoded vector of each OCR was then used as the set of features for the different machine learning classifiers. The autoencoder performance displayed on the result tables was achieved using an encoding dimension of 15.

### Sequence Wavelet Encoding

We also analyzed the fixed-window nucleotide sequences using a wavelet decomposition of the numerical representation of these sequences. Utilizing the recommended representations from Saini et al (Saini and Dewan, 2017), we created a numerical version of each nucleotide by using the electron ion interaction potential (EIIP) format, which represents the distribution of free electrons’ energies along the DNA sequence. This biologically meaningful format translates an adenine to 0.1260, guanine to 0.0806, thymine to 0.1335 and cytosine to 0.1340. The suggested wavelet configuration (“db1” mother wavelet function, level 1) for the EIIP representation produced the machine learning features from the frequency components.

### Peak Attribute Encoding

Our simplest encoding utilizes peak attributes consisting of properties that can be derived from the ATAC-seq signal itself. The features consisted of the detected peak’s width, the distance from the last ATAC-seq peak (0 if it was the first peak in each chromosome), the mean normalized number of reads, the minimum and maximum normalized read coverage, the G/C ratio, and whether this peak overlapped an annotated promoter (determined from the RefSeq (O’Leary *et al.,* 2016) release 90 gene annotations, depending on the corresponding strand coding for each gene). A fixed window of a 1kbp centered at the ATAC-seq peak’s midpoint was used for the mean, minimum and maximum normalized read coverage, as well as the GC ratio, regardless of the peak dimensions.

### Classifiers

We evaluated the classification performance of random forests (RF), AdaBoost and support vector machines (SVM). The RF classifier used 1000 estimators, no bootstraping and “Gini” criterion, as determined by hyper-parameter optimization using the individual samples. Similarly, the SVM classifier was determined to use an RBF kernel, a penalty (C) of 100 and a gamma kernel parameter of 0.01, and AdaBoost set to 400 estimators using the “SAMME.R” algorithm.

Additionally, we developed three recurrent neural network (RNN) models to classify ATAC-seq peaks, utilizing the Keras (https://keras.io) framework. Two models focused on a single source of features (i.e. sequence or signal) whereas the third utilized a combination of the two. For our sequence model, we utilized the Hill et. al. approach and mapped an input sequence of nucleotides to a sequence of vectors using an embedding layer (Hill *et al.,* 2018). This sequence of vectors was then fed into a bidirectional recurrent neural network (Fig. 3) composed of gated recurrent units (GRU) (Cho *et al.,* 2014). The outputs of both directions’ final time steps were concatenated together and then fed to an output layer for prediction. The signal-based RNN received the ATAC-seq peak signal as input to a bidirectional GRU for prediction (Fig. 4). Finally, in our hybrid model (Fig. 5, 6), we utilized both sequence embeddings and peak signal by stacking them before providing the combined input to a bidirectional GRU. A learning rate of 0.01 and a hidden layer size of 150 was determined as the optimal configuration after hyperparameter optimization. All instances of the RNN models were executed with the aid of a GPU for increased computational performance.

**Figure 3.**
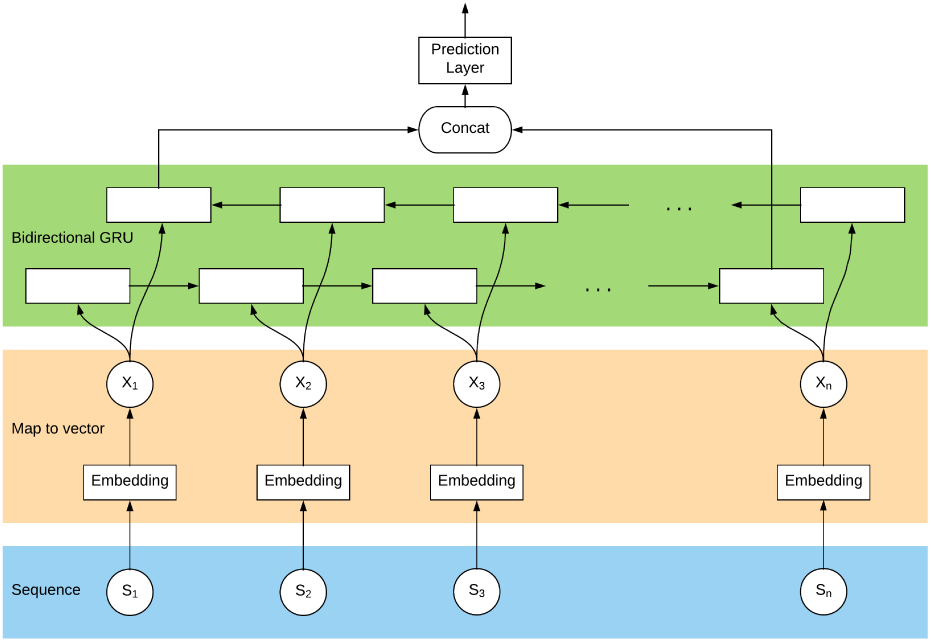
Sequence-based RNN model architecture: The nucleotides in the 1kbp evaluation window extracted from the reference genome (blue layer) were passed to an embedding layer (orange) to generate a dense vector representation from each. Each embedding vector is passed to a gated recurrent unit in each direction (green layer) to capture the long- and short-term relations between nucleotides, and the outputs from the last forward and reverse gates are concatenated to be used for the final prediction.

**Figure 4.**
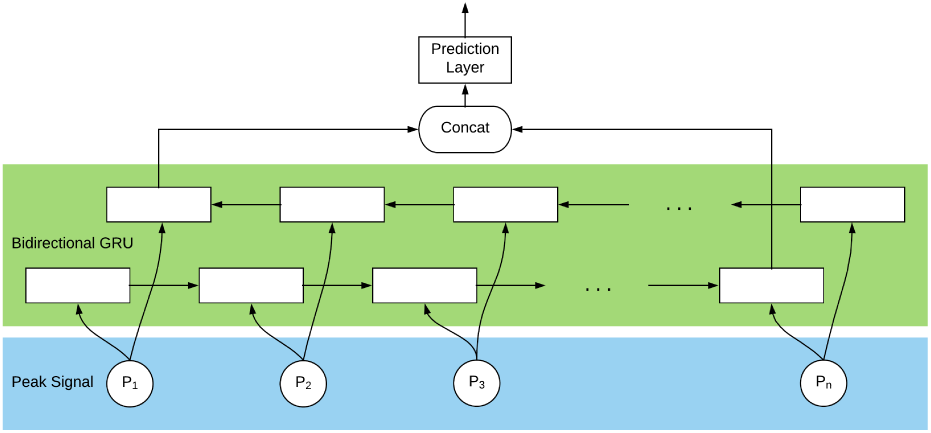
Signal-based RNN model architecture: the normalized number of reads for each nucleotide in the 1kbp evaluation window (blue layer) was passed directly to each direction of the gated recurrent unit (green layer), similarly to the sequence-based model. The output from the last gate in each direction was concatenated and used for the final prediction.

**Figure 5.**
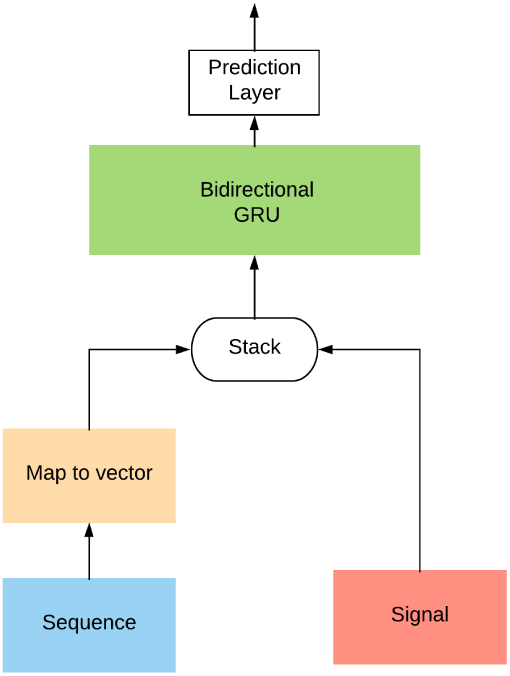
Signal/sequence hybrid model architecture: The sequence from the 1kbp evaluation window (blue) is passed to an embedding layer (yellow), to generate a dense vector representation of each nucleotide, and be combined to the vector of real values representing nucleotide-resolution normalized number of reads in that same 1kbp window (red) to create a multidimensional representation of each ATAC-seq peak. This hybrid encoding of the peak is then passed to a bidirectional gated recurrent unit, whose output is then used for label prediction.

**Figure 6.**
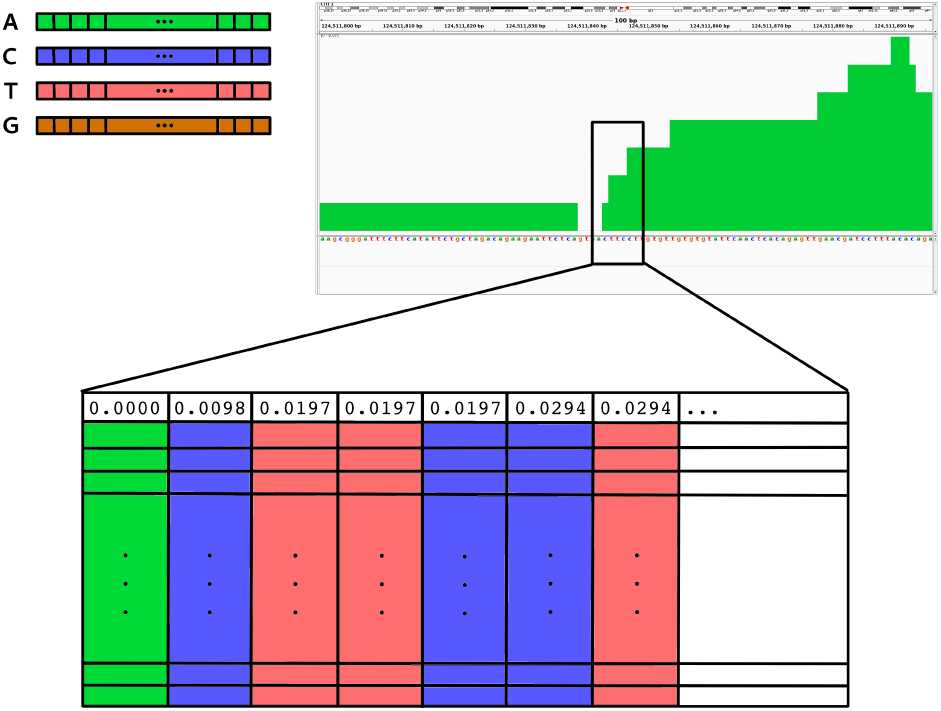
A vector embedding was trained for each nucleotide (top left, also including other base symbols following the IUPAC convention). For our signal/sequence hybrid model, we generated a multidimensional training vector for each peak combining the embedding for each nucleotide and the normalized number of reads mapped for that base. In this example, we show how a small window of this ATAC-seq region (top right) with the sequence ACTTCCT would be represented in two dimensions (bottom, one nucleotide per column), with the first row reflecting the normalized read coverage for each of those nucleotides and the rest of each column consisting of the nucleotide’s dense vector representation.

### Evaluation of models

We used F1-score (the harmonic mean between precision and recall), weighted by support, to compare the performance of the classifiers with different data encoding schemes. The F1-score for each sample was calculated using a “leave-one-out” strategy, reserving the sample in question for testing and using all others to train the machine learning classifier.

## Results

We sought to determine if we could classify ATAC-seq peaks as to whether they overlap sites of transcription initiation. We utilized nascent transcription data as our ground truth for presence of RNA polymerase initiation (e.g. bidirectional transcription) in a region. A random labeling of ATAC-seq peaks from all samples was used as a baseline, which resulted in an F1-score of 0.5 or worse, regardless of the model. The distribution of positive and negative labels for this random baseline model was matched to the observed distribution, with a probability of 0.495 for positive labels. In contrast, the classification results using labels derived from the detected bidirectionals from nascent transcription are displayed on Fig. 7. We obtained substantial improvements in F1-score over our baseline when classifying ATAC-seq regions on whether they contain RNA polymerase initiation.

**Figure 7.**
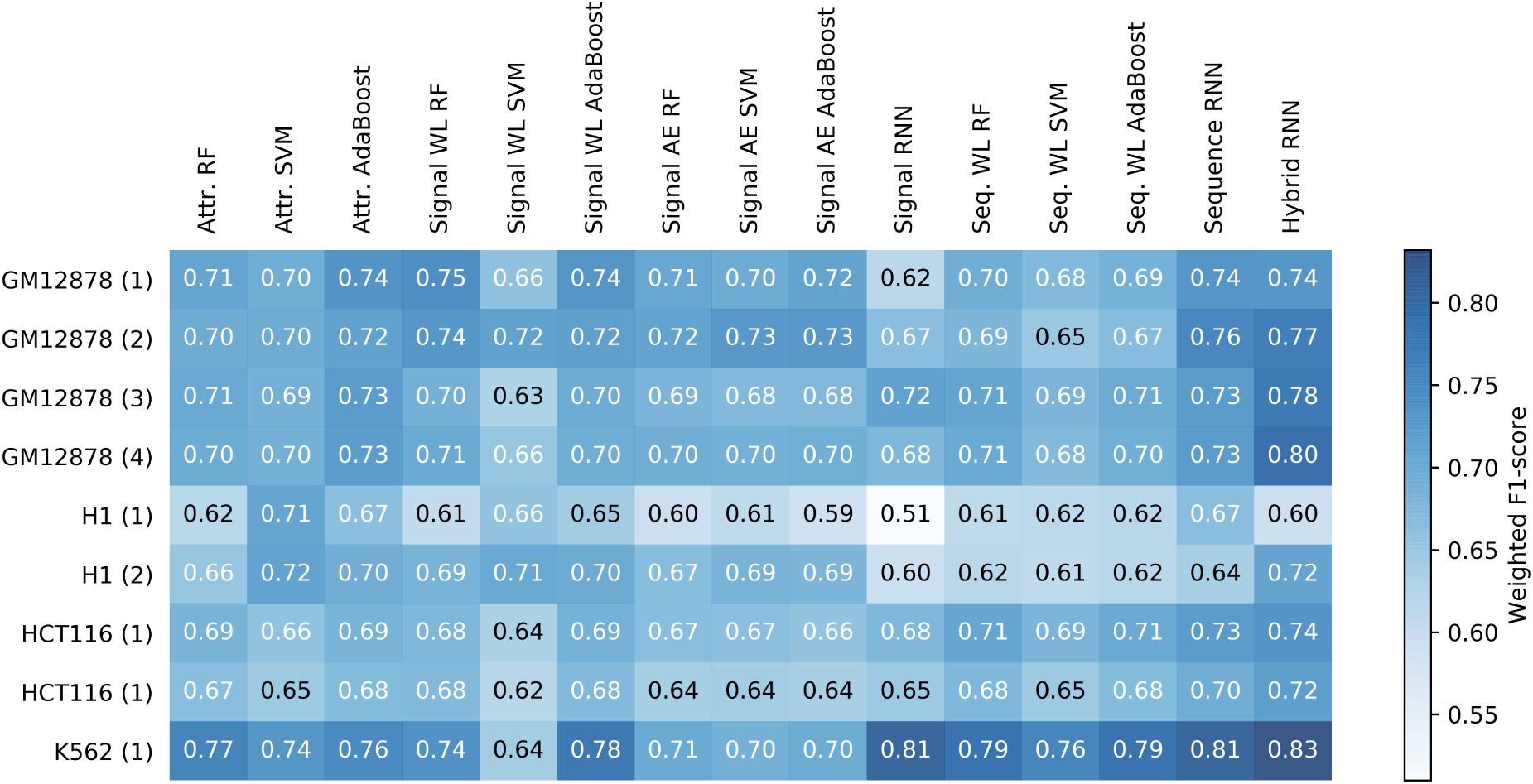
Summary of all weighted F1-scores (shown both numerically and as heatmap) obtained when attempting to detect RNA polymerase initiation (bidirectional signals in nascent transcription) from every classifier and data encoding scheme, using the sample denoting each row as test data and the rest of samples as training. Each sample was tested using multiple representations: features derived from peak attributes (Attr.), the level-1 discrete wavelet components (WL) from the peak signal itself (single nucleotide resolution), a reduced-dimension feature set from the signal vector using an AutoEncoder architecture (AE), a vector embedding of the sequence in the evaluated regions (Seq. RNN), the level-1 discrete wavelet components of the EIIP representation of this same sequence (Seq. WL), and a hybrid model that combined the sequence embeddings and the signal vector into a single training matrix per OCR. The machine learning classifiers tested were random forests (RF), support vector machines (SVM), AdaBoost and recurrent artificial neural networks (RNN).

Considering models where features are encoded as simple attributes, we note that not all features proved to be useful. A feature drop-out test, where we repeatedly attempted classification of our best-performing sample by leaving out one feature at a time, resulted in nearly identical F1-scores in the absence of the minimum number of normalized reads, the distance from the last peak, and the GC-richness of a peak’s region. The latter was not expected, as we have observed a bias in GC content for bidirectional nascent transcription (Azofeifa *et al.,* 2018). A test of feature importance using the extra trees classifier confirmed these findings (Supplemental Fig. 1). When considering sequence based features, the sequence wavelet encoding was outperformed by the RNN nucleotide vector embedding-based model.

In every scenario, the RNN-based hybrid approach that combines a vector embedding representation of the sequence on top of the signal from ATAC-seq peaks, in a multidimensional training vector per peak, outperformed all other models. This would hint that the biochemical significance of each nucleotide matters when measuring the normalized number of reads of a peak. It is worth noting, however, that some of the wavelet-based signal processing approaches resulted in a similar performance for some of the test samples, yet required significantly less computing resources. For example, an AdaBoost classifier trained on the first-level components of the discrete wavelet transform of the ATAC-seq peak signal, appeared to perform nearly a well as the RNN signal-based model, on average. The improvement in performance by the RNN became more evident when incorporating histone marks as labels, as we can see in Fig. 12.

The classification performance varied across the samples used for testing. Interestingly, the datasets sequenced to a greater depth were not necessarily the ones generally obtaining the best F1-scores, rendering this attribute orthogonal to classifier performance. However, the quality of the samples did appear to influence the models. For example, the sample “H1 (1)” (see Supplemental Table 1) was issued a warning for over-represented sequences by the FastQC (Andrews, 2010) quality control software and was the worst performing test set when using exclusively sequence-based features. This poor classification performance was reflected both on the wavelet components of the EIIP sequence representation, and the vector embedding-based representation using recurrent neural networks. The other replicate from this same dataset, which didn’t exhibit the sequence over-representation warning (H1 (2)), resulted in a better sequence-based classification performance.

Fig. 8 illustrates the overlap of true/false positives with a total of 8339 peaks contained in all datasets. The vast majority of these peaks were correctly predicted as “overlapping active transcription”, perhaps unsurprisingly since they are, by virtue of being consistent, always present in our training dataset. From the 8339 peaks contained in all datasets, very few are mis-classified as false positives by any model. Hence the majority of errors reside within the sample specific subset of peaks.

**Figure 8.**
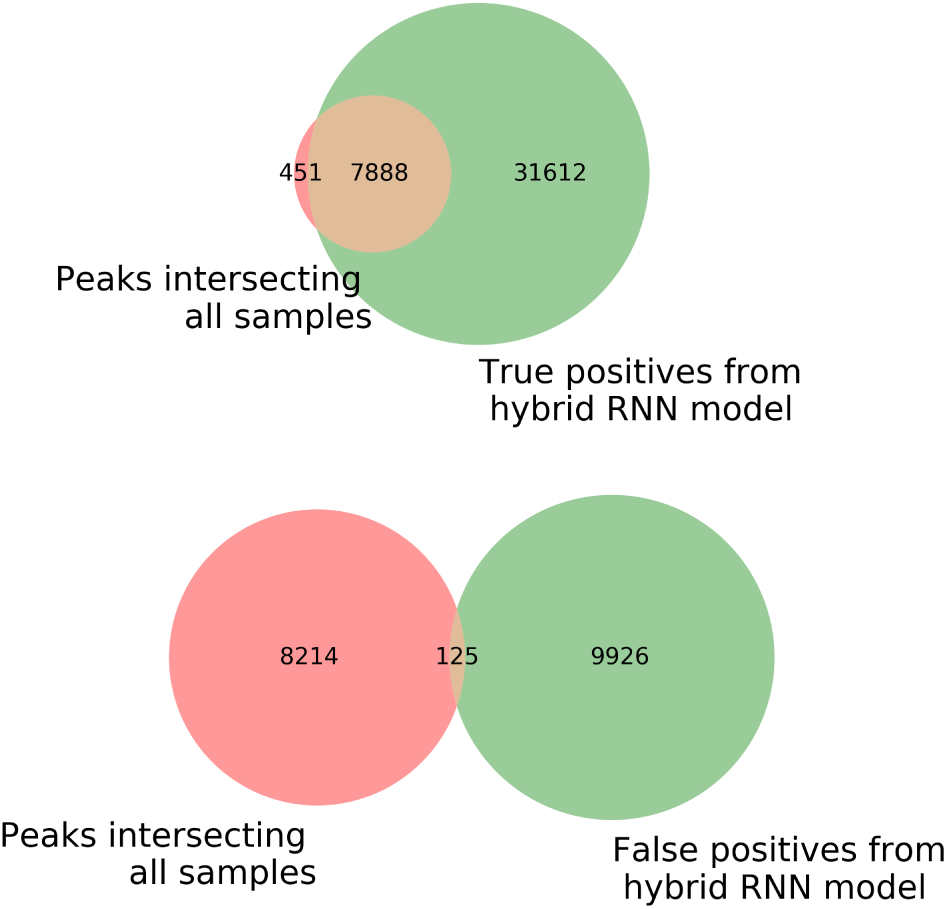
ATAC-seq peaks common to all samples (n=8339, pink circles) are overwhelmingly represented among those correctly identified by the classifiers (top green circle), while only a minority of those are among the incorrect classifications (bottom green circle). This is illustrated by the GM12878 (1) results using the hybrid RNN model true positives (n=39500) and false positives(n=10051), and is representative of all other samples regardless of the classification scheme used.

Speculating about technical ways to improve the OCR classification performance, a weighted ensemble of machine learning classifiers was an option to consider. Supplemental Fig. 2 illustrates the percentage of classification errors exclusive to three radically different models and three representative samples. When performing error analysis for the same sample across the multiple classifier types and data encoding schemes, we see that some of our false positives and many false negatives are exclusive to the specific model used. This indicates that an ensemble of machine learning classifiers may provide improve performance on any individual dataset.

To evaluate intrasample error further, we examined meta plots of false positives and false negatives within both the ATAC-seq and nascent datasets. Importantly, these analyses show bidirectional transcription signal in the nascent dataset within both error classes (Fig. 9.A). The bidirectional signal in false positives suggests these regions may be transcribed but were not called by existing bidirectional detection tools. However, given the relatively low signal signal in these regions, they are likely the difficult to call cases. Likewise, we observed that the distribution of the mean number of reads in ATAC-seq for each peak shifted towards smaller read counts for false positives, compared to true positives. This indicates that the classifiers have a tendency to mis-classify weaker peaks (those with a lower read coverage) as “not overlapping active transcription” (Fig. 10). Taken together, these results highlight the impact of sample complexity and hence strength of signal on our classification results. In contrast, false negatives show a robust bidirectional signal indicating that these regions are legitimately missed. Interestingly, these regions have a distinct shape within the ATAC-seq data (Fig. 9.B) which indicates that the ATAC-seq peaks and the bidirectional nascent transcripts may be somewhat offset from each other in these regions. Alternatively, the classifiers may be able to discriminate OCRs overlapping RNA Pol II from other types of transcription activity such as RNA Pol I or III, as observed by a recent study identifying transcription regulatory elements from nascent transcription (Wang *et al.,* 2018). It is of interest in future work to determine why these peaks are missed and whether they harbor distinct features that could be leveraged to improve our classifier.

**Figure 9.**
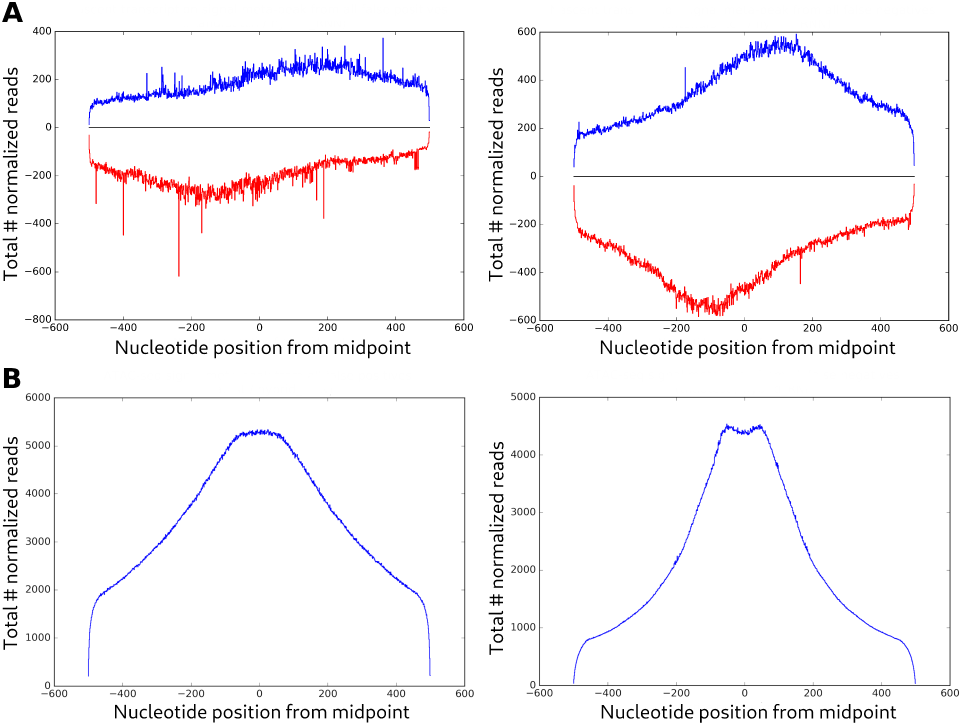
Meta-peaks for all type I (left, n=68766) and type II (right, n=87084) classification errors, across all samples, from the hybrid RNN model results. No significantly different common shape is observed in either case for nascent transcription data (A), among samples and classification schemes. The presence of bidirectional signal across so many peaks categorized as false positives may indicate that either the coverage at these regions is generally low, or that some of these may be true positives instead, and missed by the current bidirectional detection tools. The same distinct shapes were observed in each case for ATAC-seq signal (B), regardless of the model used.

**Figure 10.**
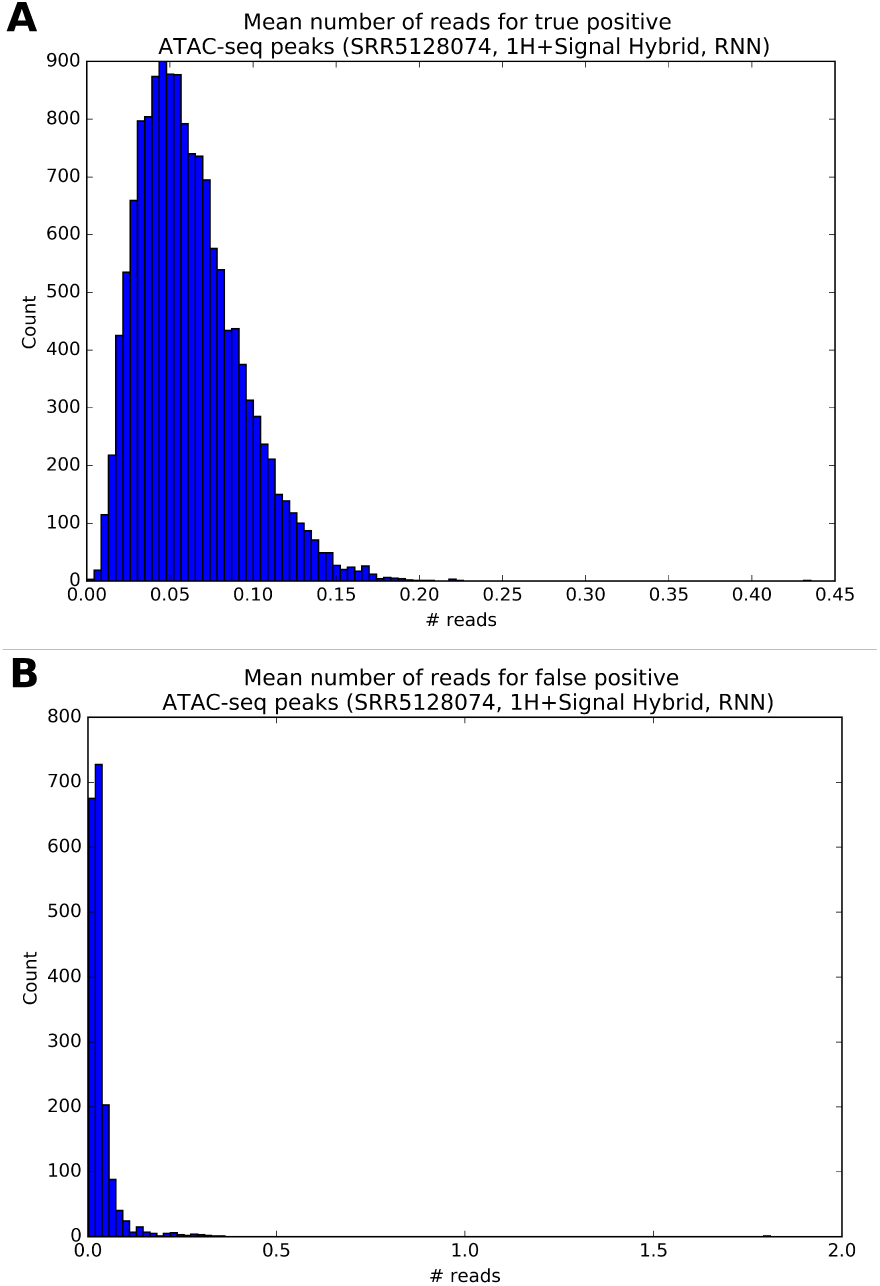
Histogram of the mean number of reads in a 1kbp window of ATAC-seq true positive peaks (A, n=8804) and false positive peaks (B, n=2038) for the K562 (1) sample utilizing the hybrid representation and RNNs, which illustrates a trend observed in all other samples and classification schemes. In general, false positives tend to be strongly represented by a lower read count.

As certain histone marks (H3K4 in particular (Core *et al.,* 2014)) correlate with transcription, we also sought to determine whether our models could be used to detect histone marks, as shown in Supplemental Fig. 3. In this case the performance is slightly better than when detecting RNA polymerase activity, perhaps in part because both histone mark ChIP-seq and ATAC-seq both measure aspects of chromatin, and thus are closely related experiments. Our results with histone marks are consistent with recent efforts to utilize ATAC-seq to identify regions labeled by ChromHMM as enhancers (Thibodeau *et al.,* 2018). However, here we infer the ChIP-seq histone modification data directly from nucleotide-resolution ATAC-seq peaks.

As histone marks indirectly imply RNA polymerase involvement, a natural next step was to ask whether we can complement the detected regions of active transcription with the histone mark ChIP-seq peaks. The combined approach did yield slightly improved performance (increased F1 score, Fig. 12) relative to the distinct individual models. These histone marks may denote regions of RNA polymerase activity that couldn’t be detected by the nascent transcription samples we used (and perhaps could be detected if sequenced at a higher depth), consistent with our false positives having relatively low but persistent bidirectional signal.

Given there is a rather large overlap between the histone marks utilized and bidirectionals in general, the improvement suggests that the two data types complement each other. The general magnitude of errors when using both data types is equivalent to using histone marks alone. Although roughly half of the errors made when training on histone marks alone were also seen with purely nascent transcription training (Fig. 11). The improved classification (Supplemental Fig. 4) reflects correctly predicting many of the peaks that were mis-classified by each labeling scheme alone. The temporal dynamics between RNA polymerase and histone marks is not entirely clear. Consequently it is interesting to speculate whether the combination of labels expands the spectrum of transcriptional activity that can be captured by ATAC-seq.

**Figure 11.**
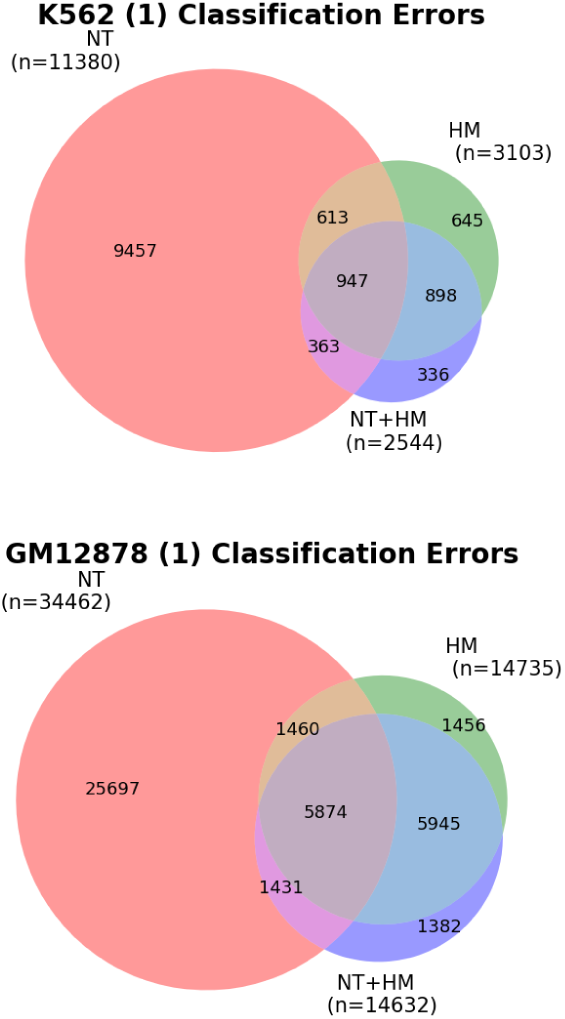
Number of overlapping peaks that resulted in an erroneous classification (combining type I and II errors) when labeling peaks from nascent transcription (NT), histone marks (HM), and the combination of both (NT+HM) for training and testing. When training on datasets with the combined labels, it appears that the classifier only misses a small subset of the same peaks from either single-assay label alternative alone. This may indicate that learning from HM patterns of signal and sequence also helps when recognizing peaks overlapping NT, and vice versa.

**Figure 12.**
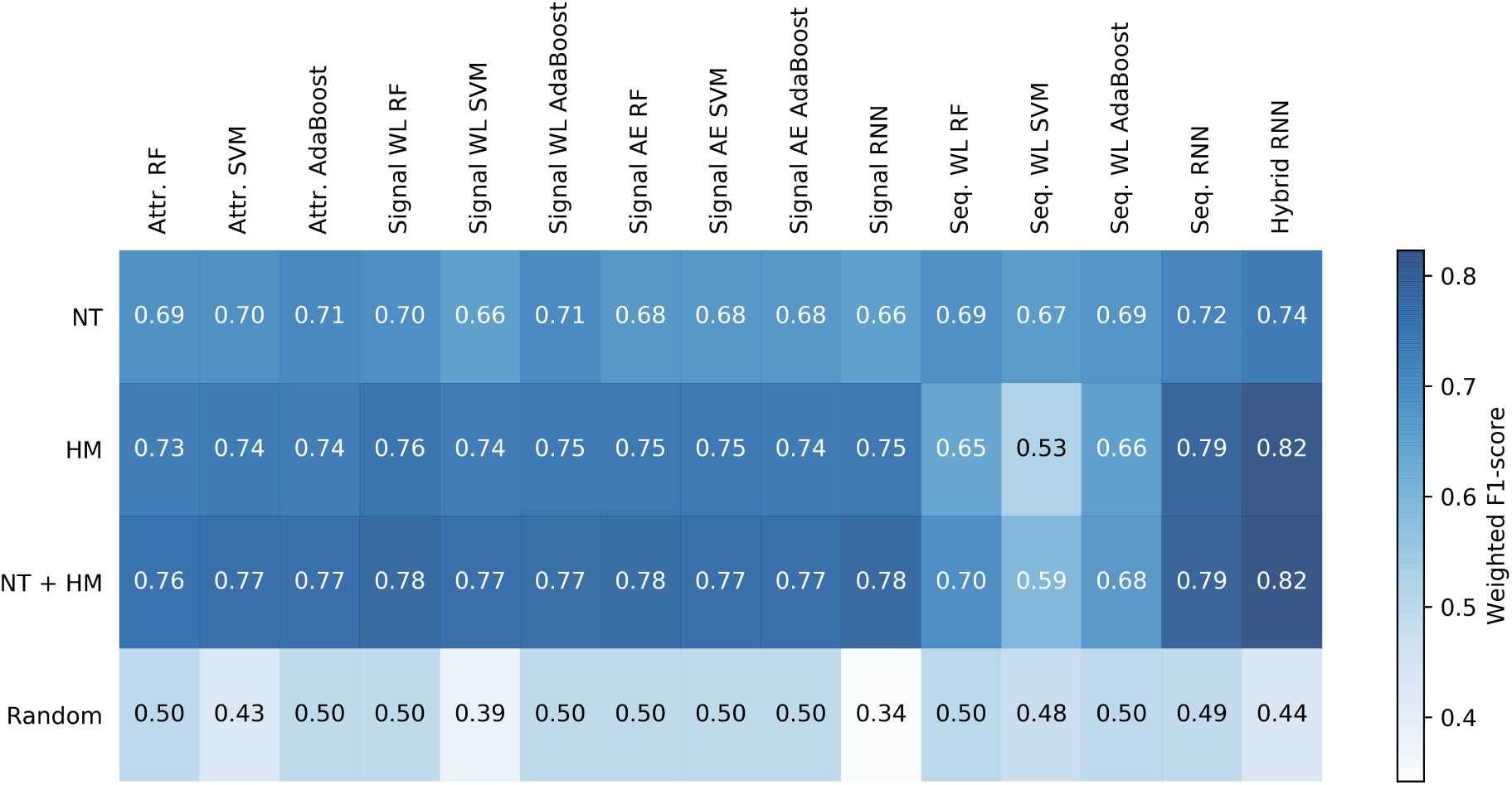
Average of F1-scores over all samples for the different models, and the different types of labels used as “ground truth”. The first row indicates the F1 averages when trying to predict RNA polymerase initiation from nascent transcription (NT) datasets, using bidirectional regions detected by Tfit and FStitch. The next row lists F1 averages detecting enhancer/promoter activity from histone marks (HM). Next, we used a combination of nascent transcription assays and histone marks, as a way to use ATAC-seq peaks to detect functional TF binding in general. Lastly, random labels matching the distribution across all samples were used as a baseline calculation.

## Discussion

Because of its relative simplicity and utility across a broad range of cell types and counts, it is advantageous to maximize the information obtained from ATAC-seq. Here we sought to determine whether we could distinguish ATAC-seq peaks that denote active transcription initiation from those that just indicate accessible chromatin. The underlying assumption was that RNA polymerase would induce particular signatures within ATAC-seq peaks. The preliminary results indicate there is promise in this approach, with the classifiers reaching a maximum F1-score of 0.83 (0.92 when also including histone marks in training labels).

The combination of sequence and signal features resulted in the best classification performance. This is perhaps unsurprising given that sequence features are necessary for regulators, such as transcription factors and yet by itself fails to provide cell type and condition specific context information. Here we utilized a diversity of cell types and sample sources, in an attempt to maximize generalizability to experiments in different cell types and from different laboratories, but possibly at the expense of classifier performance. The machine learning features from our hybrid representation of signal and sequence depicted in Fig. 6 will likely be applicable to other experimental assays and classification tasks, as well.

The fact that we can classify peaks using a signature in frequency space leads us to speculate whether this is related to the periodicity associated to nucleosome positioning, or in accessible chromatin locations. A recent study (Pich *et al.,* 2018) described in depth a periodicity in mutation rates associated to major/minor groove DNA regions, and to whether these were facing towards or away from histones. The transposase used in ATAC-seq may be able to insert in preferred locations in nucleosome protected regions as well, following minor or major groove preference patterns. Perhaps the wavelet components we capture are associated to these characteristics, besides the more obvious transposase inserts that would be mapped to linker DNA between nucleosomes.

The main ground truth label used for the classification of each ATAC-seq peak was determined by bidirectional peaks from GRO-seq or PRO-seq assays performed on the same cell line and conditions. Our error analysis of individual ATAC-seq peaks suggests that the tools utilized (Tfit (Azofeifa *et al.,* 2018) and FStitch (Azofeifa *et al.,* 2014)) miss a small subset of bidirectionals. A similar analysis using dREG (Wang *et al.,* 2018) showed similar overall results (data not shown), albeit with distinct problematic regions. While this lack of ground truth complicates, it reflects the inherent difficulty in detecting lowly transcribed bidirectionals (also known as transcriptional regulatory elements) within nascent transcription with absolute certainty. Regardless, nascent transcription assays are still ideal for labeling active RNA polymerase locations.

A possible extension of this work would be the development of an ensemble of machine learning classifiers, that combines different models (e.g. peak attributes classified by SVM, and hybrid signal/sequence features classified by RNN). This stack of learners may result in a more balanced classifier. The different types of quality control warnings could also be used to weight the different machine learning classification approaches accordingly in an ensemble setting. Another interesting next step would be to explore how tools that employ ATAC-seq data for differential analysis of TF activity such as DAStk (Tripodi *et al.,* 2018b) can benefit from using one of these classifiers as a pre-filter, by using only those peaks considered as “overlapping active Pol II recruitment”, instead of all peaks from each sample. Alternatively, DAStk could weight peaks based on their classification outcome, giving those predicted to overlap RNA polymerase larger weights.

## Acknowledgements

We would like to thank Dr. Lucas Monzon for the insightful discussions on signal processing and signature detection, and the ENCODE (Davis *et al.,* 2018) consortium and associated labs for the publicly available samples.

## Funding

This work was funded in part by the Olke C. Uhlenbeck Graduate Fellowship (IJT), a NSF ABI DBI-12624L0 (RDD) and NIH R01GM125871 (RDD, IJT). High Performance Computing resources, provided by the BioFrontiers Computing Core at the University of Colorado Boulder, funded by NIH 1S10OD012300.

## References

Alipanahi, B. et al. (2015). Predicting the sequence specificities of DNA-and RNA-binding proteins by deep learning. Nature Biotechnology, 33, 831.

Allen, M. A. et al. (2014). Global analysis of p53-regulated transcription identifies its direct targets and unexpected regulatory mechanisms. eLife, 3, e02200.

Andrews, S. (2010). FastQC: A Quality Control tool for High Throughput Sequence Data.

Azofeifa, J. et al. (2014). FStitch: A Fast and Simple Algorithm for Detecting Nascent RNA Transcripts. In Proceedings of the 5th ACM Conference on Bioinformatics, Computational Biology, and Health Informatics, BCB ’14, pages 174–183, New York, NY, USA. ACM.

Azofeifa, J. G. et al. (2018). Enhancer rna profiling predicts transcription factor activity. Genome Research, 28(3), 334–344.

Barrett, T. et al. (2013). NCBI GEO: archive for functional genomics data sets—update. Nucleic Acids Research, 41(D1), D991–D995.

Chen, Y. et al. (2016). De novo deciphering three-dimensional chromatin interaction and topological domains by wavelet transformation of epigenetic profiles. Nucleic Acids Research, 44(11), e106–e106.

Cho, K. et al. (2014). Learning phrase representations using rnn encoder-decoder for statistical machine translation. In EMNLP.

Core, L. J. et al. (2014). Analysis of nascent RNA identifies a unified architecture of initiation regions at mammalian promoters and enhancers. Nature Genetics, 46(12), 1311–1320.

Davis, C. A. et al. (2018). The Encyclopedia of DNA elements (ENCODE): data portal update. Nucleic Acids Research, 46(Database issue), D794–D801.

Di Tommaso, P. et al. (2017). Nextflow enables reproducible computational workflows. Nature Biotechnology, 35, 316–319.

Fuglerud, B. M. et al. (2017). A c-Myb mutant causes deregulated differentiation due to impaired histone binding and abrogated pioneer factor function. Nucleic Acids Research, 45(13), 7681–7696.

Fullard, J. F. et al. (2018). An atlas of chromatin accessibility in the adult human brain. Genome Research, 28(8), 1243–1252.

Graps, A. (1995). An introduction to wavelets. IEEE Computational Science and Engineering, 2(2), 50–61.

Hill, S. T. et al. (2018). A deep recurrent neural network discovers complex biological rules to decipher RNA protein-coding potential. Nucleic Acids Research, 46(16), 8105–8113.

Kelley, D. R. et al. (2016). Basset: Learning the regulatory code of the accessible genome with deep convolutional neural networks. bioRxiv.

Kelso, T. W. R. et al. (2017). Chromatin accessibility underlies synthetic lethality of SWI/SNF subunits in ARID1a-mutant cancers. eLife, 6.

Lee, G. et al. (2019). PyWavelets: Wavelet Transforms in Python. Contribute to PyWavelets/pywt development by creating an account on GitHub. original-date: 2013-07-22T20:10:04Z.

Melgar, M. F. et al. (2011). Discovery of active enhancers through bidirectional expression of short transcripts. Genome Biology, 12(11), R113.

Min, X. et al. (2017). Predicting enhancers with deep convolutional neural networks. BMC Bioinformatics, 18(13), 478.

Mitra, A. and Song, J. (2012). WaveSeq: A Novel Data-Driven Method of Detecting Histone Modification Enrichments Using Wavelets. PLOS ONE, 7(9), e45486.

Nguyen, N. et al. (2014). A wavelet-based method to exploit epigenomic language in the regulatory region. Bioinformatics, 30(7), 908–914.

O’Leary, N. A. et al. (2016). Reference sequence (RefSeq) database at NCBI: current status, taxonomic expansion, and functional annotation. Nucleic Acids Research, 44(D1), D733–745.

Pich, O. et al. (2018). Somatic and germline mutation periodicity follow the orientation of the DNA minor groove around nucleosomes. Cell, 175(4), 1074–1087.e18.

Rendeiro, A. F. et al. (2016). Chromatin accessibility maps of chronic lymphocytic leukaemia identify subtype-specific epigenome signatures and transcription regulatory networks. Nature Communications, 7, 11938.

Saini, S. and Dewan, L. (2017). Comparison of Numerical Representations of Genomic Sequences: Choosing the Best Mapping for Wavelet Analysis. International Journal of Applied and Computational Mathematics, 3(4), 2943–2958.

Schep, A. N. et al. (2015). Structured nucleosome fingerprints enable highresolution mapping of chromatin architecture within regulatory regions. Genome Research, 25(11), 1757–1770.

Setty, M. and Leslie, C. S. (2015). SeqGL Identifies Context-Dependent Binding Signals in Genome-Wide Regulatory Element Maps. PLOS Computational Biology, 11(5), e1004271.

The ENCODE Project Consortium (2012). An Integrated Encyclopedia of DNA Elements in the Human Genome. Nature, 489(7414), 57–74.

Thibodeau, A. et al. (2018). A neural network based model effectively predicts enhancers from clinical ATAC-seq samples. Scientific Reports, 8(1), 16048.

Tripodi, I. J. and Gruca, M. (2018). Nascent-Flow. https://github.com/Dowell-Lab/Nascent-Flow.

Tripodi, I. J. et al. (2018a). An ATAC-seq pipeline wrapped in NextFlow that can be run by Jupyter (ATACFlow).

Tripodi, I. J. et al. (2018b). Detecting differential transcription factor activity from ATAC-Seq data. Molecules, 23(5), 1136.

Wang, Z. et al. (2018). Identification of regulatory elements from nascent transcription using dREG. bioRxiv, page 321539.

Zhou, J. and Troyanskaya, O. G. (2015). Predicting effects of noncoding variants with deep learning-based sequence model. Nature Methods, 12, 931.

