## Supplemental Materials for "ATAC-seq signal processing and recurrent neural networks can identify RNA polymerase activity"

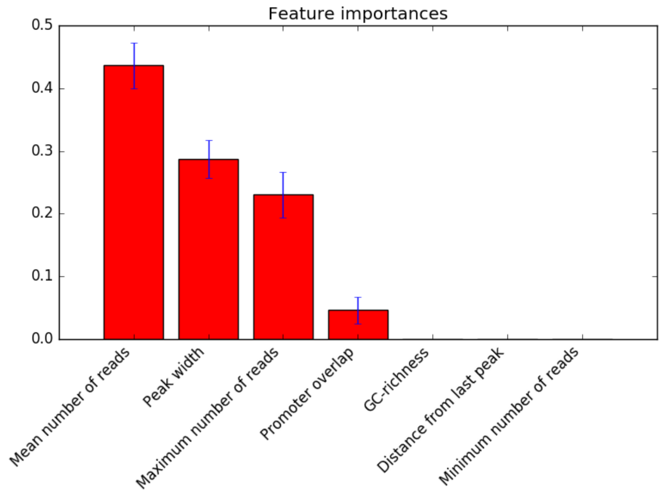


Figure 1. Importance of the peak attribute-derived features, as determined by an extra trees classifier. This result was consistent with the leave-one-out test.

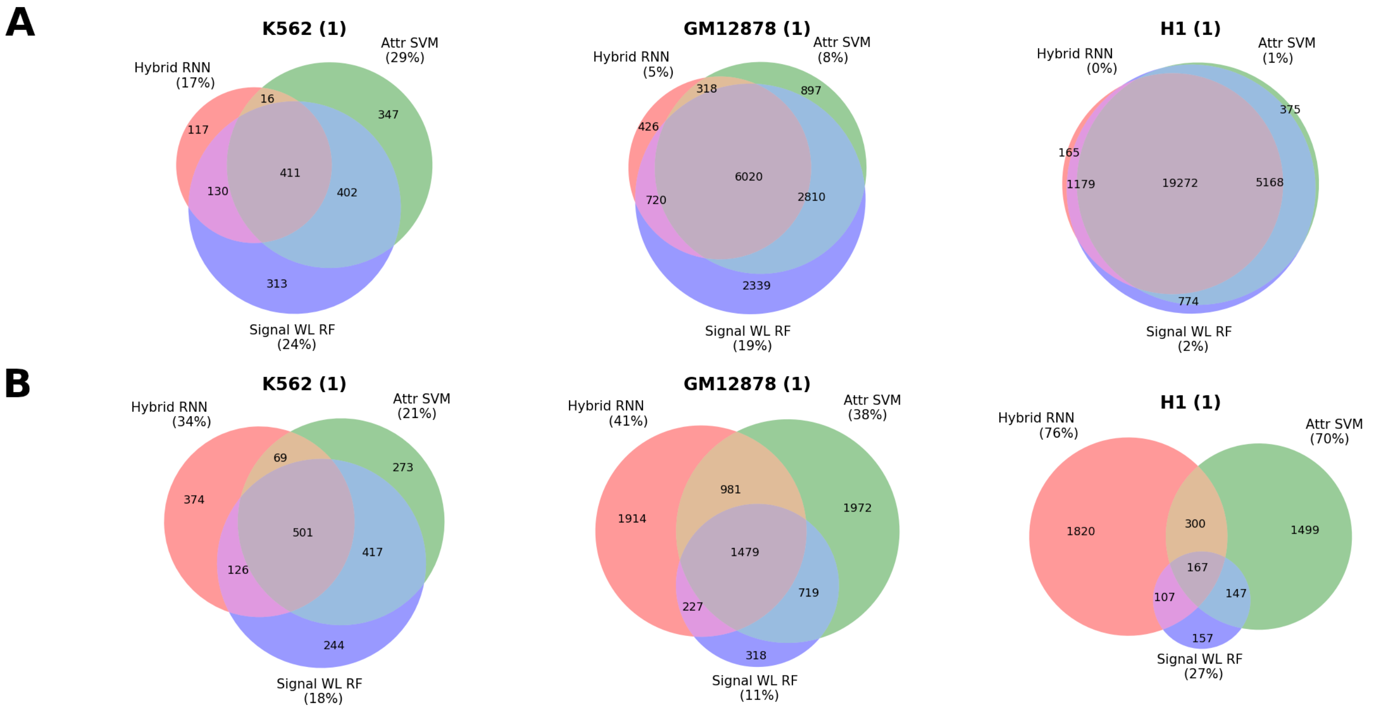


Figure 2. Overlap for false positive (**A**, n=1736, 13530, and 27090, respectively) and false negative (**B**, n=2004, 7610, and 4197, respectively) ATAC-seq peaks across the three samples (columns), between three approaches: A peak signal/sequence hybrid model using RNNs (red circle), a wavelet decomposition of the peak signal classified using random forests (purple circle), and the peak attributes classified using SVM (green circle). In each case, the number between parenthesis indicates the percentage of classification errors exclusive to that model. We note a higher number of false negative calls are unique to each approach, indicated by the smaller overlaps in **B**, hinting that an ensemble machine learning classifier approach would be worth exploring in future work.

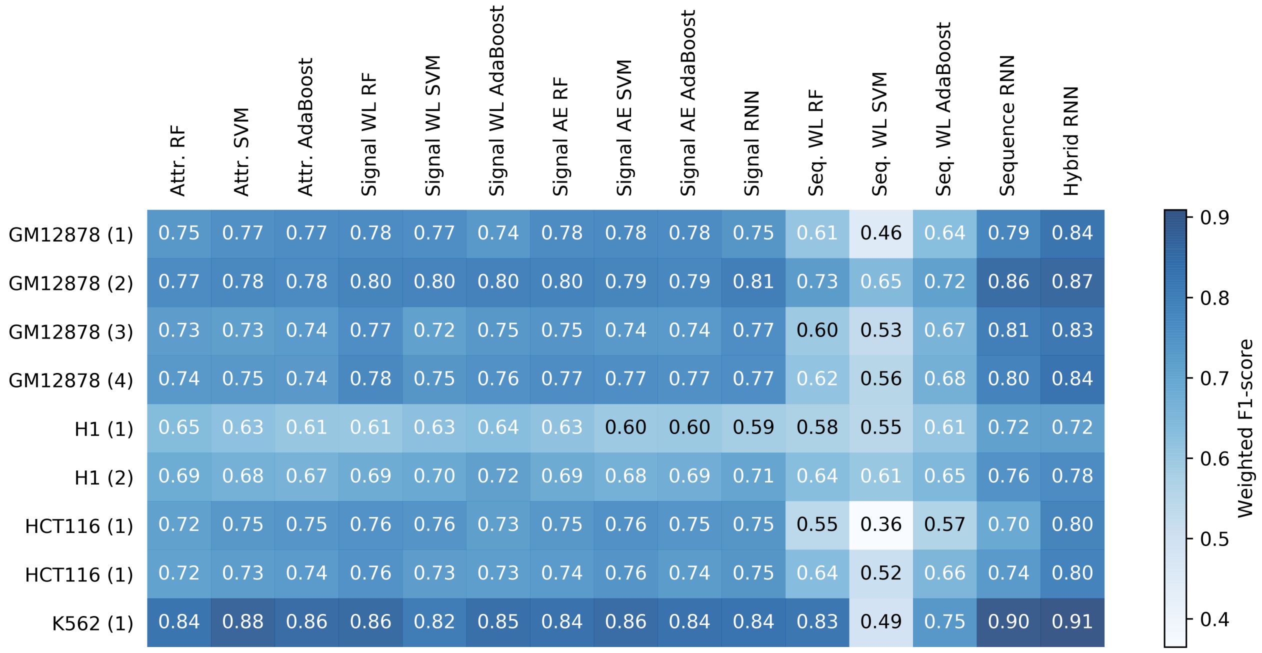


Figure 3. Summary of all weighted F1-scores obtained when attempting to detect histone marks denoting enhancers and promoters, from every classifier and data encoding scheme, using the sample denoting each row as test data and the rest of samples as training.

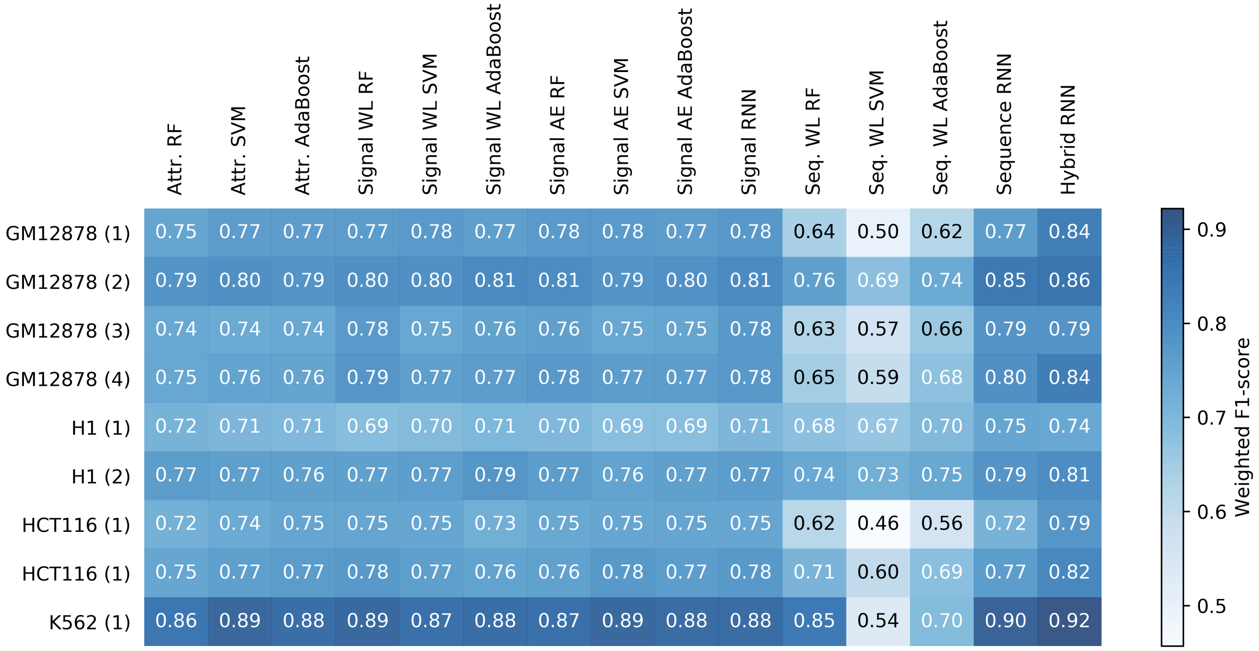


Figure 4. Summary of all weighted F1-scores obtained when attempting to detect putative functional TF binding, by combining the detected bidirectional peaks from nascent transcription with histone marks, from every classifier and data encoding scheme, using the sample denoting each row as test data and the rest of samples as training.

| Sample Name | Accession | Cell Type | FastQC Warnings | FastQC Red Flags | Nr. Reads |
|---|---|---|---|---|---|
| GM12878 (1) | SRR1822165 Schep *et al.* (2015) | Myeloid B-cell (GM12878) | Overrepresented sequences (1,2) | Per base sequence content (1,2)<br>Sequence Duplication Levels (1,2)<br>Kmer Content (1,2) | 192.9M |
| GM12878 (2) | SRR1822166 Schep *et al.* (2015) | Myeloid B-cell (GM12878) | Overrepresented sequences (1,2) | Per base sequence content (1,2)<br>Sequence Duplication Levels (1,2)<br>Kmer Content (1,2) | 56.6M |
| GM12878 (3) | SRR1822167 Schep *et al.* (2015) | Myeloid B-cell (GM12878) | Per tile sequence quality (1) | Per tile sequence quality (2)<br>Per base sequence content (1,2)<br>Sequence Duplication Levels (1,2)<br>Kmer Content (1,2) | 85.2M |
| GM12878 (4) | SRR1822168 Schep *et al.* (2015) | Myeloid B-cell (GM12878) | Per tile sequence quality (1,2) | Per base sequence content (1,2)<br>Sequence Duplication Levels (1,2)<br>Kmer Content (1,2) | 62.8M |
| H1 (1) | SRR5007258 | ESC (H1) | Per sequence GC content (1,2)<br>Sequence Length Distribution (1,2)<br>Sequence Duplication Levels (2) | Per base sequence content (1,2)<br>Sequence Duplication Levels (1)<br>Kmer Content (1,2) | 56.9M |
| H1 (2) | SRR5007259 | ESC (H1) | Per sequence GC content (1,2)<br>Sequence Length Distribution (1,2) | Per base sequence content (1,2)<br>Sequence Duplication Levels (1,2)<br>Kmer Content (1,2) | 51.4M |
| HCT116 (1) | SRR5876158 Kelso *et al.* (2017) | Colon epithelium (HCT116) | Overrepresented sequences (2) | Per base sequence content (1,2)<br>Sequence Duplication Levels (1,2)<br>Kmer Content (1,2) | 44.7M |
| HCT116 (2) | SRR5876159 Kelso *et al.* (2017) | Colon epithelium (HCT116) | | Per base sequence content (1,2)<br>Sequence Duplication Levels (1,2)<br>Kmer Content (1,2) | 31.7M |

| | | | | | |
|---|---|---|---|---|---|
| K562 (1) | SRR5128074 Fuglerud *et al.* (2017) | Lymphoblast (K562) | Per base sequence content (1)<br>Overrepresented sequences (1,2) | Per base sequence content (2)<br>Sequence Duplication Levels (1,2)<br>Kmer Content (1,2) | 36.8M |

Table 1: Sample of the public ATAC-seq samples used for this study. For the FastQC results, the (1) or (2) indicates which of the paired-end reads displayed this warning/error.

| **Sample** | **Cell Type** | **Sample Provenance** |
|---|---|---|
| SRR1552482 | GM12878 | Core, 2014 Core *et al.* (2014) |
| SRR1552483 | GM12878 | Core, 2014 Core *et al.* (2014) |
| SRR1552485 | GM12878 | Core, 2014 Core *et al.* (2014) |
| SRR1745515 | H1 | Estaras, 2015 Estars *et al.* (2015) |
| SRR1745516 | H1 | Estaras, 2015 Estars *et al.* (2015) |
| SRR1745523 | H1 | Estaras, 2015 Estars *et al.* (2015) |
| SRR1745524 | H1 | Estaras, 2015 Estars *et al.* (2015) |
| SRR1745527 | H1 | Estaras, 2015 Estars *et al.* (2015) |
| SRR1745528 | H1 | Estaras, 2015 Estars *et al.* (2015) |
| SRR574824 | H1 | Sigova, 2013 Sigova *et al.* (2013) |
| SRR574825 | H1 | Sigova, 2013 Sigova *et al.* (2013) |
| SRR574826 | H1 | Sigova, 2013 Sigova *et al.* (2013) |
| SRR1105736 | HCT116 | Allen, 2014 Allen *et al.* (2014) |
| SRR1105737 | HCT116 | Allen, 2014 Allen *et al.* (2014) |
| SRR1224573 | HCT116 | Allen, 2014 Allen *et al.* (2014) |
| SRR2084584 | HCT116 | Chen, 2015 Chen *et al.* (2016) |
| SRR2084585 | HCT116 | Chen, 2015 Chen *et al.* (2016) |
| SRR2084586 | HCT116 | Chen, 2015 Chen *et al.* (2016) |
| SRR2084587 | HCT116 | Chen, 2015 Chen *et al.* (2016) |
| SRR2084588 | HCT116 | Chen, 2015 Chen *et al.* (2016) |
| SRR2084589 | HCT116 | Chen, 2015 Chen *et al.* (2016) |
| SRR2084590 | HCT116 | Chen, 2015 Chen *et al.* (2016) |
| SRR2084591 | HCT116 | Chen, 2015 Chen *et al.* (2016) |
| SRR828695 | HCT116 | Galbraith, 2013 Galbraith *et al.* (2013) |
| SRR828696 | HCT116 | Galbraith, 2013 Galbraith *et al.* (2013) |
| SRR828729 | HCT116 | Galbraith, 2013 Galbraith *et al.* (2013) |
| SRR1552480 | K562 | Core, 2014 Core *et al.* (2014) |
| SRR1552481 | K562 | Core, 2014 Core *et al.* (2014) |
| SRR1552484 | K562 | Core, 2014 Core *et al.* (2014) |
| SRR1554311 | K562 | Core, 2014 Core *et al.* (2014) |
| SRR1554312 | K562 | Core, 2014 Core *et al.* (2014) |
| SRR1823901 | K562 | Niskanen, 2015 Niskanen *et al.* (2015) |
| SRR1823902 | K562 | Niskanen, 2015 Niskanen *et al.* (2015) |

Table 2: Public nascent transcription samples used with Tfit and FStitch.

| **Sample** | **Cell Type** | **Sample Provenance** |
|---|---|---|
| ENCFF835NLI | GM12878 | GSM733771, Bernstein 2011 |
| ENCFF710GQV | GM12878 | GSM733772, Bernstein 2011 |
| ENCFF772RNW | GM12878 | GSM733769, Bernstein 2011 |
| ENCFF797IEH | GM12878 | GSM733677, Bernstein 2011 |
| ENCFF774QTB | GM12878 | GSM733642, Bernstein 2011 |
| ENCFF608EGM | GM12878 | GSM945188, Stamatoyannopoulos 2011 |
| ENCFF067WBB | H1 | GSM733718, Bernstein 2011 |
| ENCFF483GVK | H1 | GSE96392, Bernstein 2016 |
| ENCFF835TGA | H1 | GSM605312/GSM466739, Ren 2013 |

| | | |
|---|---|---|
| ENCFF551VUC | H1 | GSM602260/GSM602261, Ren 2013 |
| ENCFF931KXH | H1 | GSM434785/GSM605323, Ren 2013 |
| ENCFF665UWA | H1 | GSM605329/GSM789284, Ren 2013 |
| ENCFF246BAX | HCT116 | GSE95958, Bernstein 2017 |
| ENCFF786ORW | HCT116 | GSE96381, Bernstein 2017 |
| ENCFF366QBO | HCT116 | GSE95916, Bernstein 2016 |
| ENCFF403LQH | HCT116 | GSE96185, Bernstein 2017 |
| ENCFF450AGJ | HCT116 | GSM945853, Farnham 2012 |
| ENCFF004ZFO | HCT116 | GSM945304, Stamatoyannopoulos 2011 |
| ENCFF437DPT | K562 | GSM733656, Bernstein 2011 |
| ENCFF104FIT | K562 | GSM733651, Bernstein 2011 |
| ENCFF999RPS | K562 | GSE96303, Bernstein 2016 |
| ENCFF854AMS | K562 | GSM788082, Farnham 2011 |
| ENCFF501DZN | K562 | GSM788085, Farnham 2011 |

Table 3: Public ChIP-seq peaks from ENCODEDavis *et al.* (2018) samples capturing the various histone marks.

### References


Allen, M. A. *et al.* (2014). Global analysis of p53-regulated transcription identifies its direct targets and unexpected regulatory mechanisms. *eLife*, **3**, e02200.

Chen, Y. *et al.* (2016). De novo deciphering three-dimensional chromatin interaction and topological domains by wavelet transformation of epigenetic profiles. *Nucleic Acids Research*, **44**(11), e106–e106.

Core, L. J. *et al.* (2014). Analysis of nascent RNA identifies a unified architecture of initiation regions at mammalian promoters and enhancers. *Nature Genetics*, **46**(12), 1311–1320.

Davis, C. A. *et al.* (2018). The Encyclopedia of DNA elements (ENCODE): data portal update. *Nucleic Acids Research*, **46**(Database issue), D794–D801.

Estars, C. *et al.* (2015). SMADs and YAP compete to control elongation of $\beta$-catenin:LEF-1-recruited RNAPII during hESC differentiation. *Molecular Cell*, **58**(5), 780–793.

Fuglerud, B. M. *et al.* (2017). A c-Myb mutant causes deregulated differentiation due to impaired histone binding and abrogated pioneer factor function. *Nucleic Acids Research*, **45**(13), 7681–7696.

Galbraith, M. D. *et al.* (2013). HIF1a employs CDK8-mediator to stimulate RNAPII elongation in response to hypoxia. *Cell*, **153**(6), 1327–1339.

Kelso, T. W. R. *et al.* (2017). Chromatin accessibility underlies synthetic lethality of SWI/SNF subunits in ARID1a-mutant cancers. *eLife*, **6**.

Niskanen, E. A. *et al.* (2015). Global SUMOylation on active chromatin is an acute heat stress response restricting transcription. *Genome Biology*, **16**, 153.

Schep, A. N. *et al.* (2015). Structured nucleosome fingerprints enable high-resolution mapping of chromatin architecture within regulatory regions. *Genome Research*, **25**(11), 1757–1770.

Sigova, A. A. *et al.* (2013). Divergent transcription of long noncoding RNA/mRNA gene pairs in embryonic stem cells. *Proceedings of the National Academy of Sciences of the United States of America*, **110**(8), 2876–2881.
